# Areas Important for Ecological Connectivity Throughout Canada

**DOI:** 10.1101/2021.12.14.472649

**Authors:** Richard Pither, Paul O’Brien, Angela Brennan, Kristen Hirsh-Pearson, Jeff Bowman

## Abstract

Governments around the world have acknowledged the importance of conserving ecological connectivity to help reverse the decline of biodiversity. In this study we employed recent methodological developments in circuit theory to conduct the first pan-Canadian analysis of multi-species connectivity for all terrestrial regions of the country, at a spatial grain sufficient to support local land-management decisions. We developed a movement cost surface with a limited number of thematic categories using the most recently updated land cover data available for the country. We divided the country into 17 tiles and used a wall-to-wall, omnidirectional mode of Circuitscape on each tile in order to assess ecological connectivity throughout entire landscapes as opposed to strictly among protected areas. The resulting raw current density map of Canada revealed heterogenous patterns of current density across the country, strongly influenced by geography, natural barriers, and human development. We included a validation analysis of the output current density map with independent wildlife data from across the country and found that mammal and herpetofauna locations were predicted by areas of high current density. We believe our current density map can be used to identify areas important for connectivity throughout Canada and thereby contribute to efforts to conserve biodiversity.

## INTRODUCTION

In a 2019 global assessment, the Intergovernmental Science-Policy Platform on Biodiversity and Ecosystem Services^1^ warned that as many as one million species face extinction unless action is taken to address the underlying causes of biodiversity loss. Habitat loss is one of the main drivers behind the decline in biodiversity^2,3^ and its effect can be further exacerbated when remnant habitat patches are fragmented and separated by barriers that impede the movement of wildlife among the patches^4,5^. Consequently, reversing the decline of biodiversity will require not only protecting more natural habitat but also maintaining and restoring ecological connectivity in the wider landscape to maintain the flow of individuals, genes and ecological processes^1^. This will be particularly important in the face of climate change, in order to facilitate species’ range shifts^6,7^.

Governments around the world have acknowledged the importance of ecological connectivity. In 2020, the Parties to the Convention on Migratory Species (CMS) noted that the declines in ecological connectivity and habitat quality have contributed to a worsening of the conservation status of migratory species and affirmed that maintaining and restoring ecological connectivity is one of their top priorities^8^. The following year, the United Nations General Assembly issued Resolution 75/271 that stressed the need for cooperation “on the enhancement of connectivity between ecosystems and cooperation in order to maintain healthy and intact ecosystems and habitats, which are needed to conserve biodiversity…”.

To meet international and domestic expectations, land-use planners and managers need to be able to objectively define and map areas important for connectivity. This can be challenging because connectivity, or the “degree to which the landscape facilitates or impedes movement among resource patches”^9^ is species-dependent, since species typically differ in their movement traits and resource needs. Consequently, areas that are important for one species may not be for others^10^. Nevertheless, large-scale connectivity initiatives are quite likely to require the implementation of a multi-species approach due to the typical goals of land-use planning. Inevitably, this requires collapsing information from multiple species into a single model somewhere in the work flow^11^. Some have used the approach of empirically parameterizing resistance surfaces and then combining the resulting connectivity maps for a handful of species^12–14^. Doing so, however, still requires selecting which species to include and species-specific data, which is often limited in availability and quality. Furthermore, combining such maps can lead to the multiplication of errors and result in higher levels of uncertainty in the final product^15^. Another approach is to generate a single connectivity map for multiple species based on the assumption that the movement of many species is impeded by unnatural, human-modified land cover features such as cities and roads^16,17^. For example, Koen et al.^18^ created a single, generalized movement cost layer reflecting the assumed permeability of the landscape for several species and were able to simultaneously predict areas of high functional connectivity for one large mustelid species and several small amphibian and reptile species.

The growing interest in ecological connectivity has led to the development of several models and software for assessing connectivity^19,20^. One of the more commonly used methods in recent years involves electrical circuit theory^21,22^, especially since the introduction of the software package Circuitscape in 2008^23–25^. In circuit theory models, animal movement or gene flow mimic the flow of electricity through circuits and allow for multiple movement paths with varying levels of resistance. Unlike least-cost paths, the methodology underpinning circuit theory is analogous to a random walk approach that assumes no *a priori* knowledge of the landscape^26^. A circuit theory model can produce a current density map, reflecting the probability of an animal moving across any point within a landscape as it moves from source to destination. Importantly, maps produced using circuit theory models have proven to be accurate in identifying areas important for connectivity^14,20,27^.

In recent years there have been several research developments and methodological improvements to circuit theory that have facilitated its use for connectivity mapping over large areas. The first applications of circuit theory tended to connect source and destination locations (hereafter referred to as nodes) to measure the cost of travelling between these nodes (effective resistance), and to provide a map of current density reflecting the most probable travel paths^24,26^. Subsequent research identified that map edges can lead to biased effective resistance estimates^28^ and node placement can lead to biased current density estimates^18^. Consequently, it has been recommended that it is better to place nodes on the outside edge of a buffer placed around the area of interest if the objective is to predict connectivity across landscapes in general as opposed to among specific locations^18^.

It has also been shown that an omnidirectional current density map can be produced using methods of placing nodes outside of the study area and allowing current to flow in all directions across the map^18,29,30^. An omnidirectional connectivity map predicts the probability of use across the entire modelled landscape, independent of source and destination nodes, which is valuable when the source and destination of all movement is unknown and highly relevant for generalized land use planning activities^10^. More recently, a moving window version of omnidirectional connectivity has become available in the Omniscape package^31^. Phillips et al.^20^ compared the implementation and results of the three common omnidirectional connectivity methods (wall-to-wall, point-based, and Omniscape) and found that all three methods produced very similar current density patterns, while the wall-to-wall method was the fastest for computer processing.

Researchers have also assessed how some aspects of the cost maps used as circuit theory inputs can affect connectivity assessments. For example, current density measures have been found to be sensitive to landscape definition, including spatial grain, thematic resolution, and the number of geospatial layers^32^. Furthermore, the size of the map tile used in the analysis can influence patterns of current density, with larger tiles reducing this effect^33^. In addition, current density estimates have been found to be relatively insensitive to absolute cost values in the inputted cost surface, provided that the rank order of the costs is accurate^34^. Taken together, we consider that a cost surface for omnidirectional connectivity mapping should have few, large tiles, and it should have relatively few categories (themes) that can be accurately measured, and where relative costs can be accurately assessed.

Canada is a member state of the UN, a party to the Convention on Biological Diversity (CBD), and the second largest country in the world. To date, the only national-scale analysis of ecological connectivity of Canada was restricted to the country’s forested regions using a resistance layer with a low thematic resolution of forest vs non-forest^35^. Although that study was able to use a resistance surface with a fine spatial grain (25 x 25 m), the study had to divide Canada into relatively small tiles (25 x 25 km) due to computational limitations at the time. At the continental-scale, Carroll et al.^7^ identified areas important for connectivity between current climate zones and their future analogs under various climate change projections in North America, including Canada. More recently, Barnett and Belotte^36^ modelled connectivity across North America, but only among mostly large protected areas and at a coarse spatial grain (5 x 5 km). While mapping connectivity among existing protected areas is important, in many countries they represent only a small portion of the landscape^37^. In fact, only 12.5% of Canada’s terrestrial areas are currently protected^38^, less than half of the 30% target expected in the CBD Post-2020 Framework^39^.

In this study we employed recent methodological developments in circuit theory analyses to conduct the first pan-Canadian analysis of multi-species connectivity for all terrestrial regions of the country, at a spatial grain sufficient to support local land-management decisions. We developed a movement cost surface with a limited number of thematic categories using the most recently updated land cover data. Taking advantage of the developments that significantly improved the processing speed for circuit theory analyses^40^, we were able to run wall-to-wall analyses on very large tiles, using nodes in buffers outside the areas of interest. Rather than analyzing connectivity among the limited number of protected areas in Canada, we conducted an omnidirectional connectivity analysis for terrestrial landscapes, in which the potential contribution of all landscape elements to connectivity were considered and where source and destination nodes were independent of land tenure. Finally, we validated our output current density map with independent wildlife data from across the country.

## METHODS

### MOVEMENT COST SURFACE

Analyzing connectivity using circuit theory requires a cost surface reflecting the estimated cost of movement for animals across different landcover types^26,41^. Much like Koen et al.^18^, we modelled terrestrial fauna that use and move across natural land cover more successfully than through anthropogenic land cover types. This corresponds to an upstream connectivity modelling approach that integrates the needs of multiple species at the beginning of the analysis, as described by Wood et al.^11^.

We developed a movement cost surface for Canada by combining landcover layers from 16 sources. Eight of those layers were from the Canadian Human Footprint (CHF)^42^ and included built environments, nighttime lights, croplands, pasturelands, dams and reservoirs, mining, oil and gas, and forestry areas. We used a recently developed national road layer^43^ that included resource-access roads, along with a national railway layer. In addition to human modified land cover features, we included natural features considered to inhibit the movement of animals, namely elevation and slope, glaciers, lakes, and rivers^10,44-47^. We also included a layer for permanent sea ice because it is known to facilitate connectivity among islands in the Canadian Arctic archipelago for some mammal species^48^. All input layers were rasterized and/or resampled to a resolution of 300 x 300m, to match the resolution of layers from the CHF. Further details, including data sources, are available in the Supplementary Information (Supplementary Table 1).

We were interested in using current density maps from circuit theory as our main model output. Bowman et al.^34^ demonstrated that such maps are generally insensitive to the absolute value of the cost weights for land cover types so long as the relative ranking of the types are maintained (e.g., cities > crop lands > pasture lands > natural habitat). To ensure this remained true for our study area, we tested the sensitivity of current density outputs with ten cost scenarios (Table S2) using two regions of Canada (southern British Columbia - B.C., and the east coast provinces of New Brunswick, Prince Edward Island, and Nova Scotia). We used Spearman rank correlations to compare the current density values generated in the ten scenarios for 1000 randomly selected cells in each region. We then selected the cost weight scenario that was most strongly correlated with all the others for use in the subsequent analyses.

Given that we were interested in modelling connectivity for terrestrial animals that are likely to use and move more successfully through natural land cover types^49–51^, we assigned a high movement cost to anthropogenic land cover types (e.g., cities, major highways), a medium-high cost to human-modified features (e.g., agricultural lands, minor highways), a medium-low cost to more permeable human-modified land cover types (e.g., pasture land, resource roads, harvested forests) along with permanent sea ice, and a low cost to all other pixels, which we assume to be natural and more frequently used for movement. We included lakes greater than 10 hectares and rivers with flows above 28 m^3^/s in the high-cost category, consistent with similar studies and the understanding that many species avoid crossing large water bodies^10,52,53^. All 16 land cover layers were reclassified using the movement cost scheme and combined to produce a Canada-wide cost surface where each grid cell was assigned the highest cost from among the 16 layers (Supplementary Table 2).

To increase computational efficiency, we divided Canada into a series of overlapping tiles that were analysed individually and then stitched together to produce a national current density map. Knowing that we would likely be using tiles of different sizes because of the shape of the country, and knowing that tile size can affect current density estimates^33^, we conducted a sensitivity analysis comparing current density across tiles of varying sizes. We tested ten tiles of sizes ranging from 25 x 25 to 1500 x 1500 cells. To control for effects related to the composition and spatial distribution of cost values, we simulated simplistic landscapes in which all tiles were split in two, with 50% of the cells assigned a high cost and 50% assigned a low cost. We then calculated and compared the minimum, maximum, mean and standard deviation of the current densities for each tile size to identify the size above which variation in current density estimates was minimal. We then used tile sizes above that threshold but still with a reasonably efficient run time with our computing resources. Tiles were also sized and placed to include overlap with a neighbouring tile and a buffer. Following Koen et al.^18^, we used a buffer width equivalent to 20% of the average length of the sides of the tile of interest. This included buffers into the United States of America. We used individual layers from the Global Human Footprint^54^ to inform our cost layer in the U.S. portion and then used the same process described earlier to generate the cost values.

### CIRCUIT THEORY ANALYSIS

We used the wall-to-wall omnidirectional, Julia implementation, advanced mode of Circuitscape^55^ to run the circuit theory models on each tile following methods outlined by Phillips et al.^20^. The wall-to-wall method uses thin, one-pixel wide source and ground strips placed along opposite sides of a tile. Circuitscape was run in each of the four cardinal directions (east to west, west to east, north to south, and south to north) for a total of four runs per tile. The four directional runs were then combined by taking the mean of all four to produce an omnidirectional current density map for a given tile.

All current density maps were combined into a single map after their buffers were removed. For the overlapping areas of adjacent tiles, we used the mean current density value from the two areas. In a few areas however, the current density values from the two maps were different enough to create what appeared as seams at boundaries (i.e., high current density on one map meeting lower current density on the other). To address those anomalies, Circuitscape was rerun on smaller tiles (1,200 km^2^ plus buffers) centred over those seams. The new maps were then combined with the others by taking a mean to produce a single, seamless, national current density map.

### VALIDATIONS

We used independent wildlife datasets from across Canada to validate our current density map. This included GPS-collar data accessed through Movebank.org for moose (*Alces alces*)^56,57^, wolf (*Canis lupus*)^58^, elk (*Cervus canadensis*)^59,60^, and caribou (*Rangifer tarandus*)^61^ located in British Columbia and Alberta, as well as roadkill data for herpetofauna in southern Ontario and moose in New Brunswick (see Supplementary Figure 1 for general locations).

For the GPS-collar data from western Canada, we compared current density values extracted from observed locations (i.e., locations used by collared individuals) to current density values extracted from the same number of available, but unused locations. For the observed locations, we randomly selected 10 percent of each species dataset, to avoid issues associated with autocorrelation. To obtain the available, but unused locations, we used the adehabitatHR package in the R statistical computing environment version 3.6.3 to first generate a 95% utilization distribution (UD) of each full GPS dataset and then to calculate the maximum displacement distance observed in these datasets. We then buffered each UD by the corresponding maximum displacement distance and randomly selected points within the buffered area. Finally, we used t-tests and Cohen’s Effect Size d to compare the mean current densities for used and available points.

The herpetofauna road-kill data from Ontario contained 4496 locations of 17 species killed along roadways in southern Ontario (south of the French River) between 2010 and 2019. We used t-tests and Cohen’s Effect Size d to compare the mean current densities of the road-kill locations to densities found within an equal number of random locations along the same roads. For each location, we used the mean of the current densities of the focal pixel and within a 300m buffer on each side to capture current density adjacent to the roads. For the moose data in New Brunswick, we used road-kill observations only from highways with no wildlife fencing or passages to avoid biasing the analysis. Even so, moose road kills were prevalent, with varying numbers of road kills occurring in most 300m cells. Moose are very large and not likely to be missed when dead on the side of the road and so we assumed that we had an accurate count of moose road kills. Consequently, we used a Spearman rank correlation to compare the abundance of moose road kills to the mean current densities within sections of roads (focal pixel with 300m buffers as before).

## RESULTS

Using two of our tiles, we found that the current densities generated using ten cost scenarios were highly correlated (mean correlation of 0.82 for the eastern provinces and 0.88 for southern British Columbia, Supplemental Figures 2). Two of the cost scenarios (scenarios 9 and 10, Supplementary Table 2, Supplementary Figure 3) were equally the most strongly correlated with all other scenarios. Scenario 9 had costs of 1, 10, 100, and 1000, whereas scenario 10 had costs of 10, 100, 1000, and 10000. Given the similarity of the outputs from these two scenarios, we chose to proceed with scenario 9 because a low cost of 1 has been more conventionally used in previous studies^52,62^ (see Supplemental Figure 4 for final cost map).

The sensitivity analysis of the tile sizes revealed that variations in current density estimates became negligible above sizes of 150,000 km^2^ (1.7 million pixels; Supplemental Figure 5). Computational limitations meant that the largest tile we could efficiently analyse was 3,600,000 km^2^ (40 million pixels). This required us to divide Canada into 17 square tiles for the initial analysis. Addressing the anomalies at some of the seams required analysis of another 5 smaller tiles, for a total of 22 tiles (Supplemental Figure 6).

The raw current density map of Canada (Figure 1) revealed heterogenous patterns of current density across the country, strongly influenced by geography, natural barriers, and human development. Large corridors of high current density are evident in many regions, especially those proximate to bodies of water or other large barriers. For example, large areas of high current density occurred just south of James Bay, across Baffin Island, and across the Chignecto Isthmus between New Brunswick and Nova Scotia.

**Figure 1.**
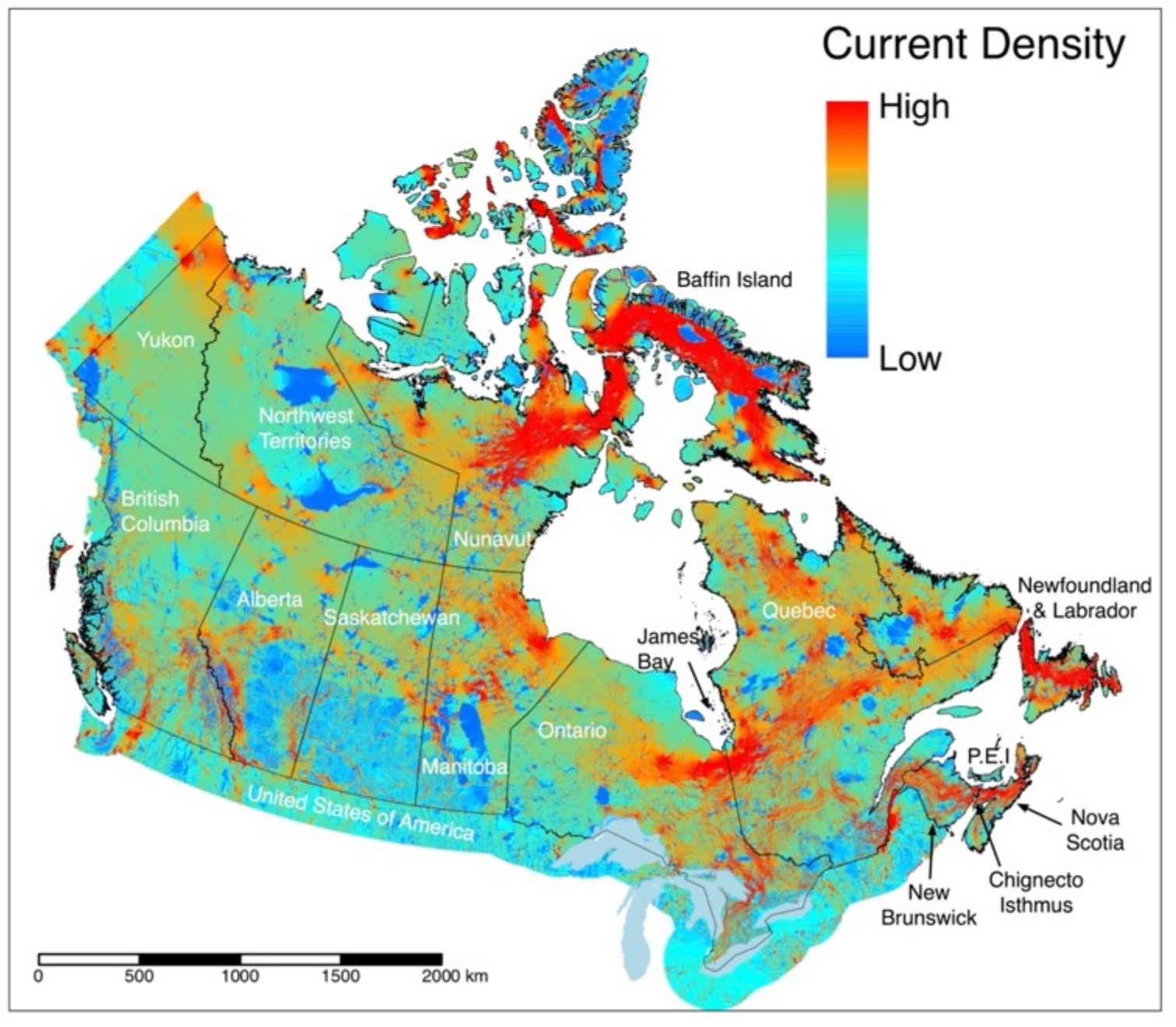
Raw current density of Canada. High current density (amperes) indicates a high probability of movement by terrestrial animals that use natural cover.

In southern Canada, where the vast majority of species at risk reside^63^, patterns of current density are difficult to see at the scale of the national map (Figure 2). However, when viewed at finer resolutions (Figures 3a, b, c), areas important for connectivity can be seen in regions considered priorities for biodiversity even within landscapes dominated by human activity. In southern British Columbia (Figure 3a) where populated areas are found in the valleys such as the Okanagan, options for movement remain in the upland resource management areas and higher mountain ranges, as reflected by diffuse patterns of current flow. Extensive agriculture throughout the prairies (Figure 3b), in contrast, leads to concentrations of high current densities following land less suitable to crops, including pasture lands used for cattle grazing. In southern Ontario (Figure 3c), Lake Simcoe is a prominent feature that leads to a north-south corridor of high current density along the shore of Georgian Bay.

**Figure 2.**
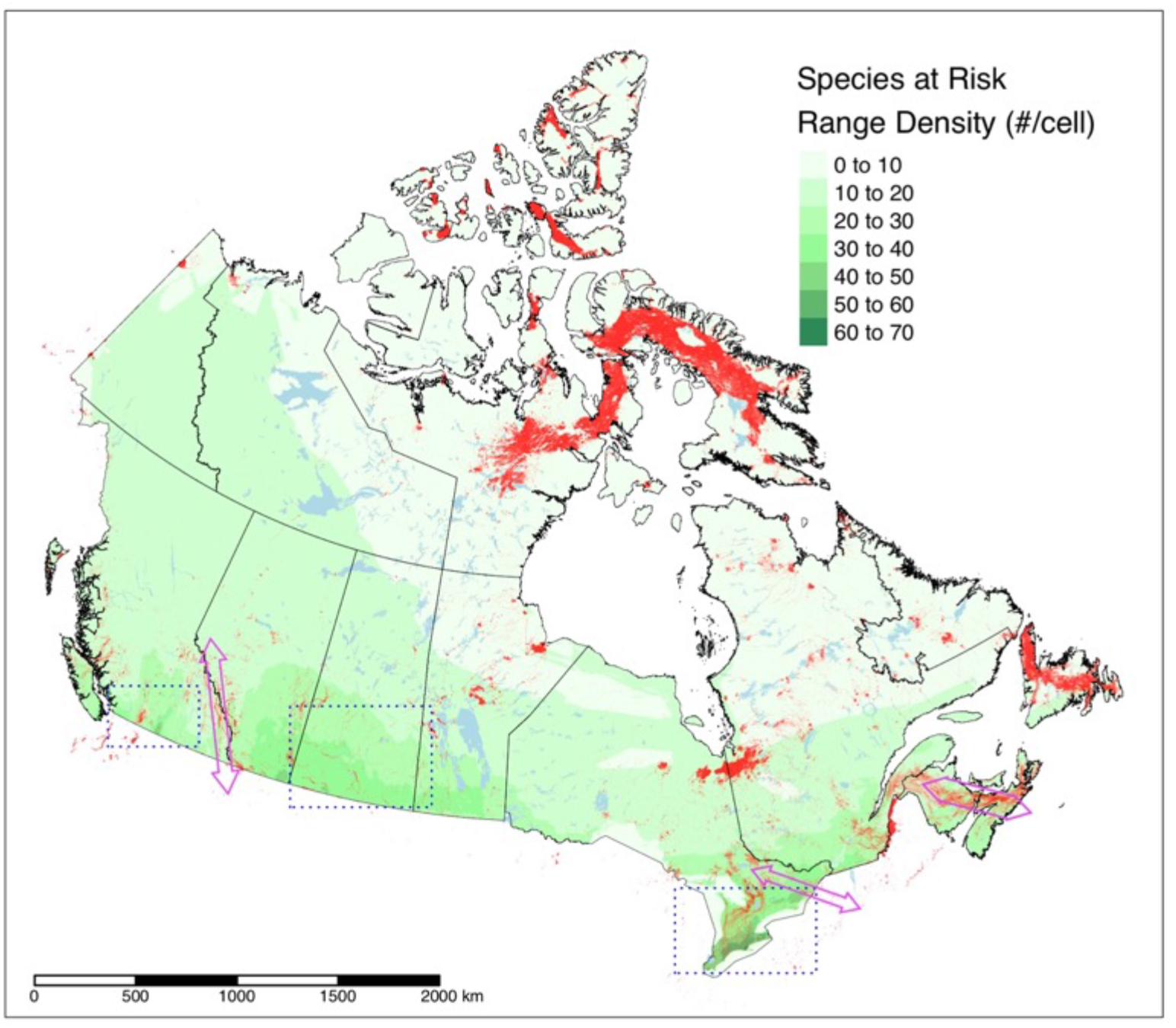
95^th^ percentile of current densities over the density of ranges for species at risk in Canada^63^. The three dashed boxes indicate the locations of the vignettes (Figures 3 a, b, & c) and the purple arrows provide examples of regions with existing connectivity conservation initiatives (Yellowstone to Yukon, Algonquin to Adirondacks, and the Chignetco Isthmus, from left to right).

### VIGNETTES

**Figure 3.**
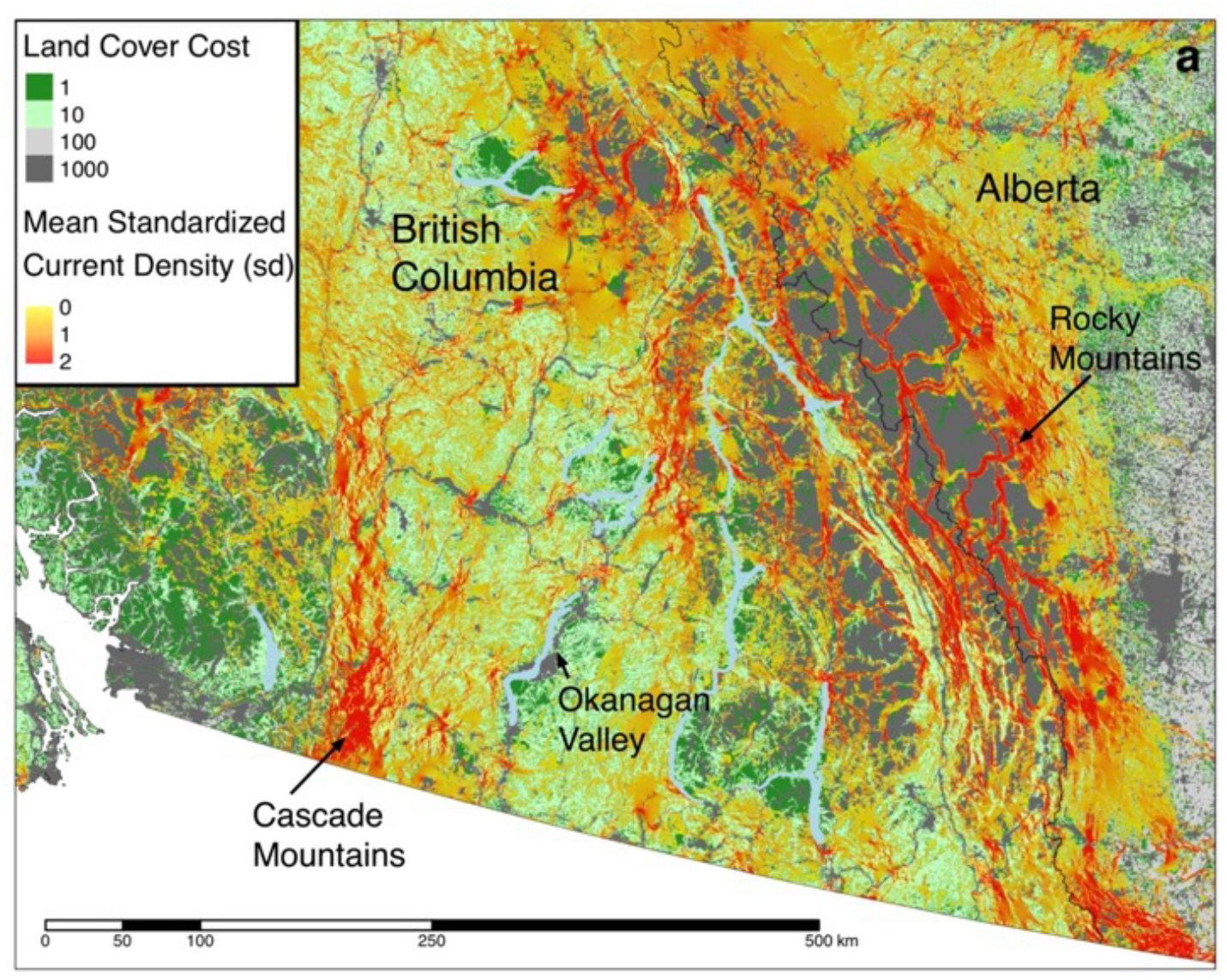
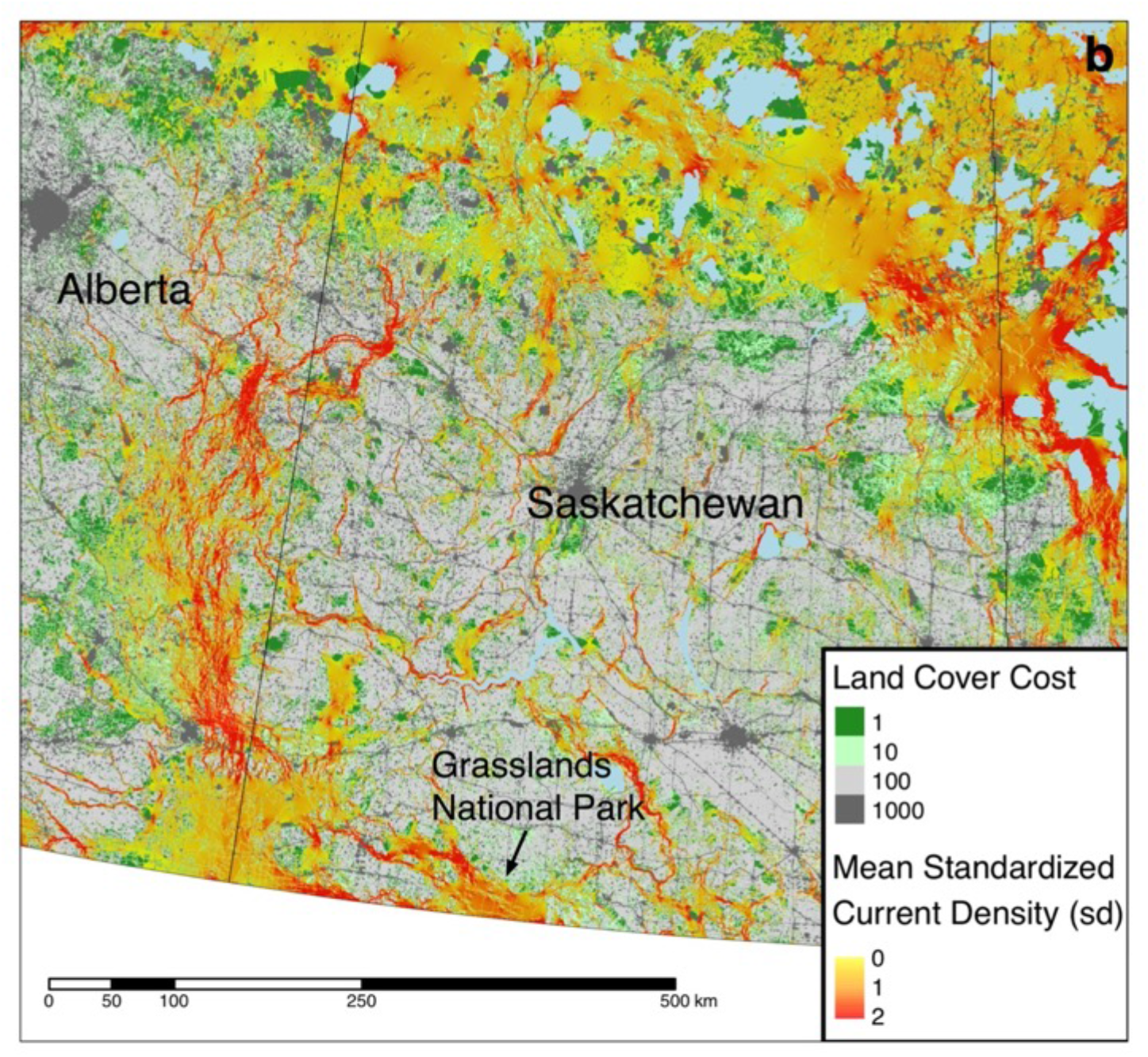
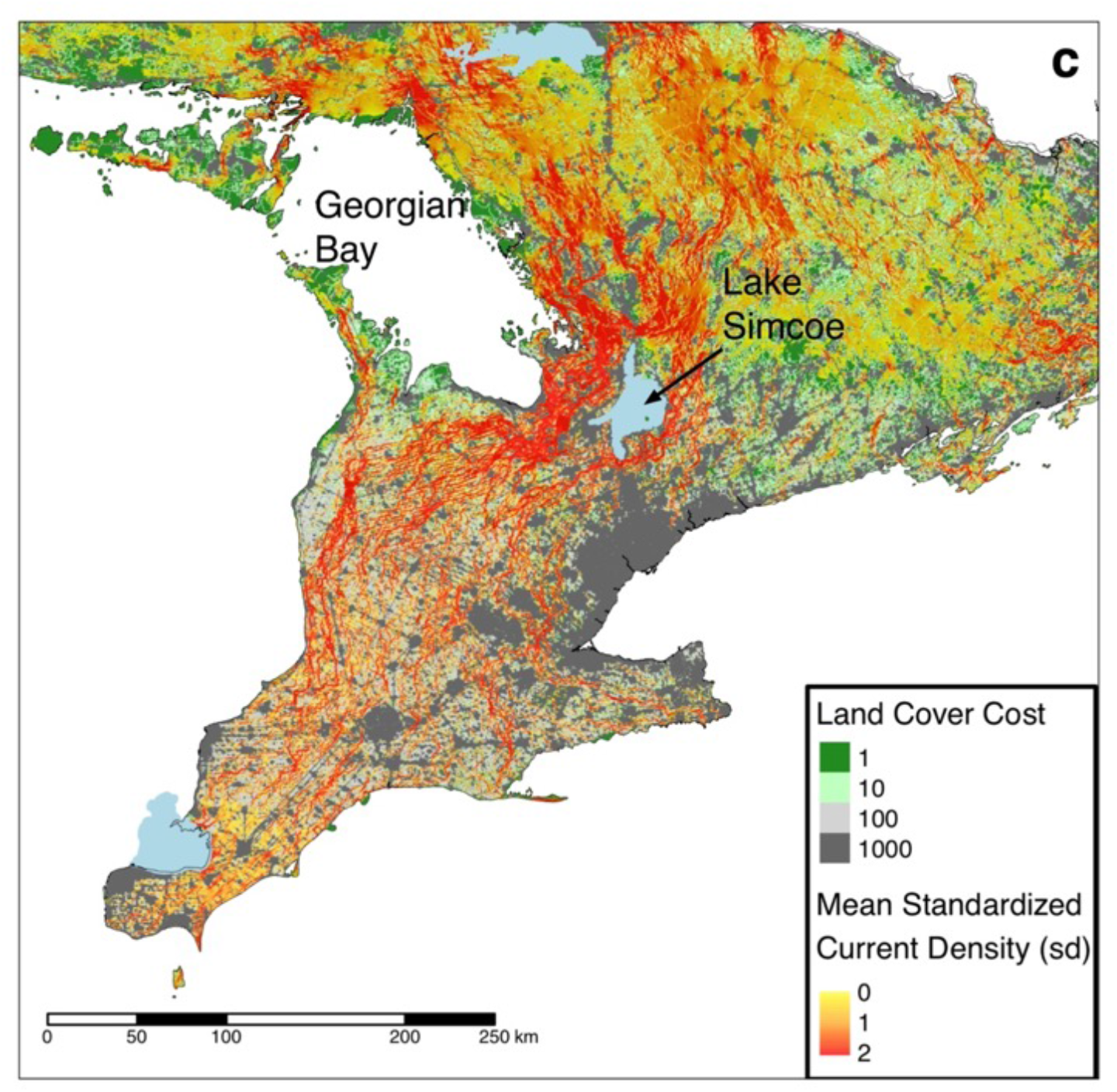
Vignettes of three priority regions for species at risk and biodiversity in Canada: a) southern British Columbia, b) southern Saskatchewan, and c) southern Ontario.

### VALIDATION RESULTS

Validation with the GPS-collar data provided evidence that mammals preferred to use areas with higher current density, as compared to current densities found in the randomly-selected, available but unused locations (Figure 4; wolf: n = 17,444, t = 69.66, P < 0.0001, d = −0.75; caribou: n = 24,546, t = 27.42, P < 0.0001, d = −0.25; moose: n = 5,518, t = 24.53, P < 0.0001, d = −0.47; elk: n = 158,543, t = 186.44, P < 0.0001, d = −0.66). Similarly, herpetofauna road kill were found more often in locations with higher current densities compared to random locations along the same roads (Figure 4; n = 4496, t = 2.04, P = 0.04, d = 0.04). There was also a positive correlation between the density of moose road kills in New Brunswick and current densities in the immediate area (Figure 5; n = 1216, ρ = 0.14, P < 0.001).

**Figure 4.**
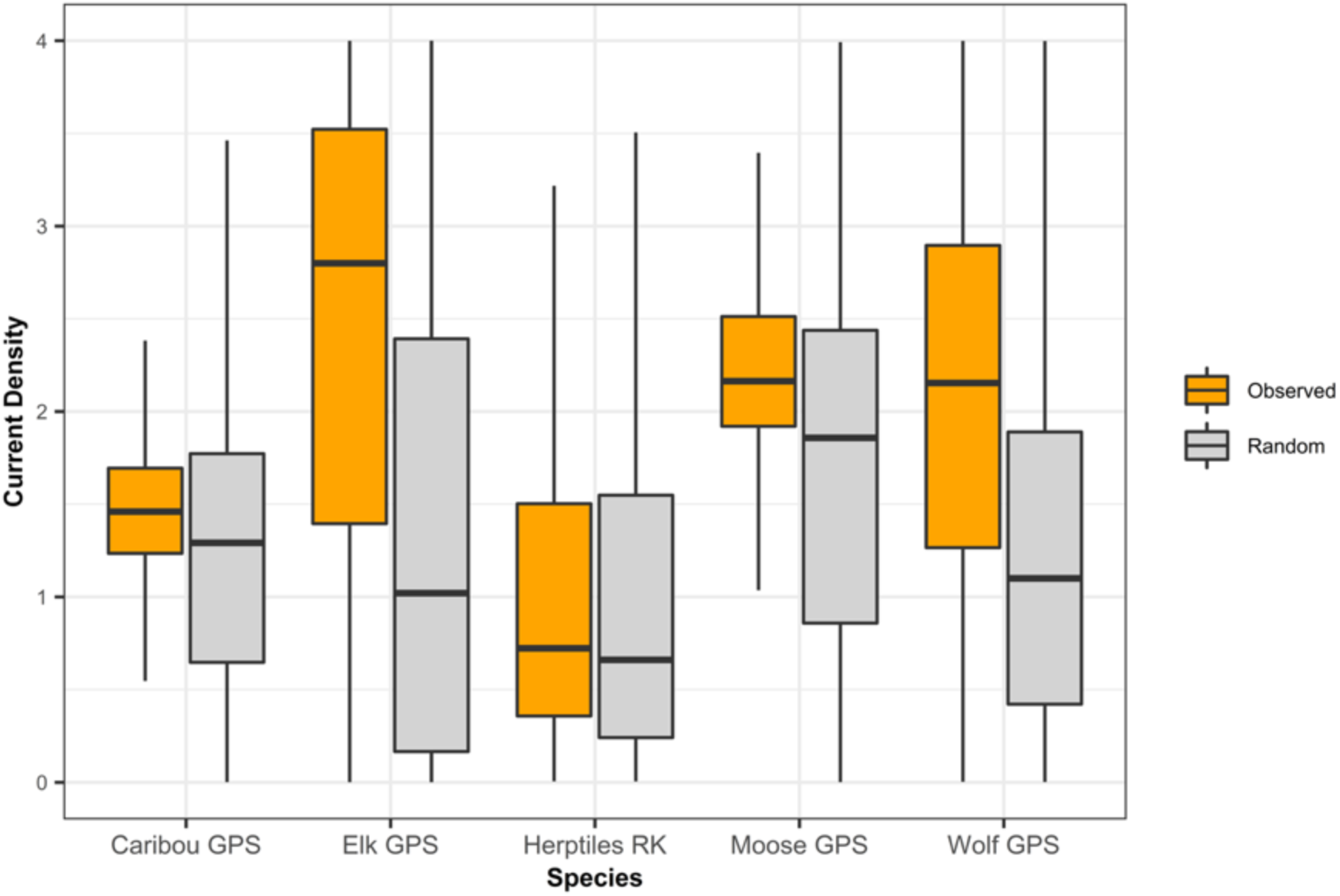
Boxplots depicting median current density (thick horizontal line) and central 50% of current density values (boxes) for wildlife locations and random locations.

**Figure 5.**
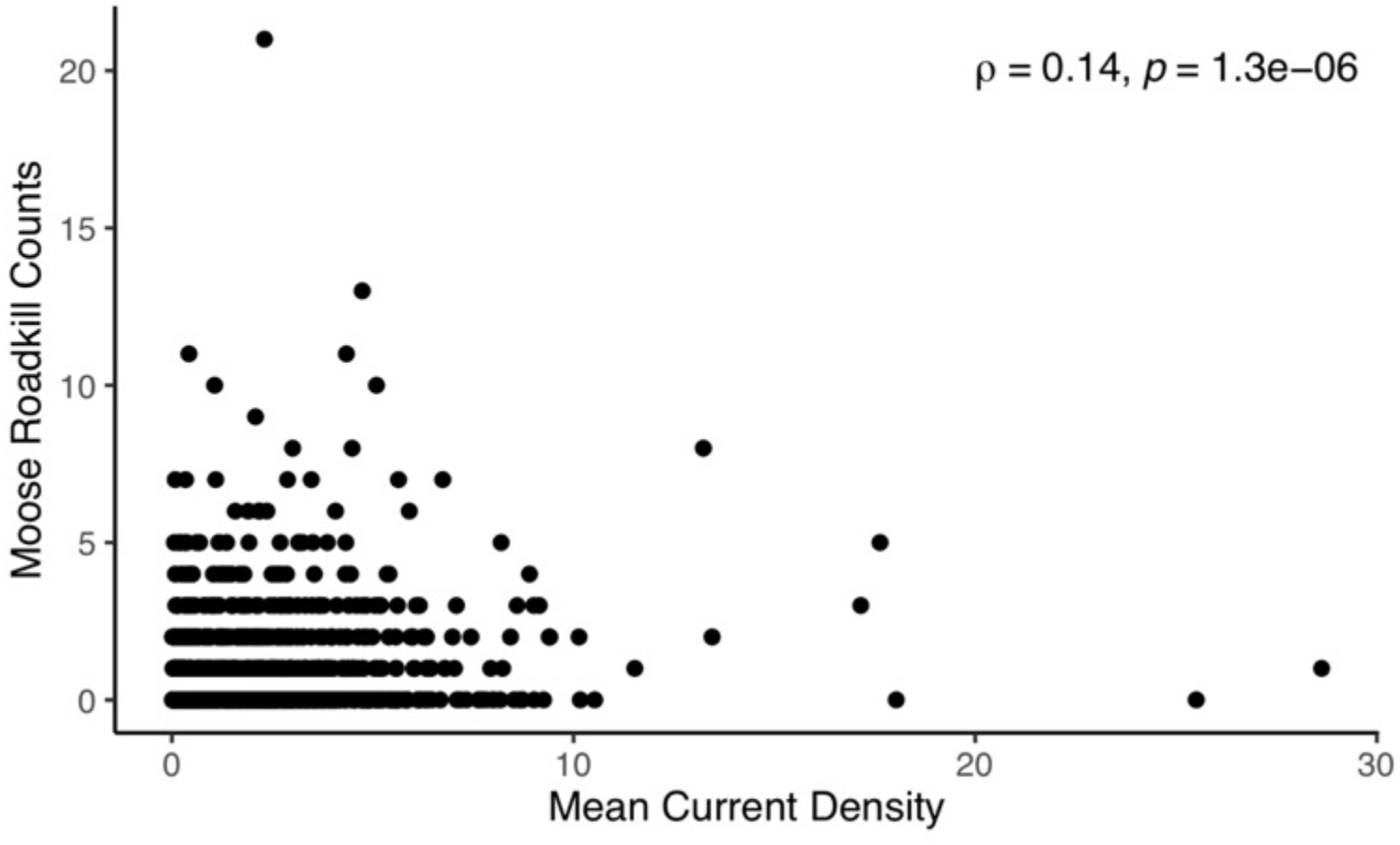
Relationship between number of moose killed on roads and mean current density in New Brunswick.

## DISCUSSION

We have produced the first map of multi-species connectivity spanning the entire country of Canada at a spatial resolution sufficient to support land-management decisions at both national and local scales. Our analysis brought together several methodological and technological advances in the application of circuit theory to measure connectivity and our current density map was validated with some, but not all, independent wildlife data. We consider our map could help Canada meet its international commitments to conserve biodiversity and establish networks of connected protected and conserved areas. This includes identifying areas that will be critical for facilitating species’ range shifts in response to climate change.

Our current density map (Figure 1) clearly shows the effects of both natural barriers and human development, highlighting areas that have a high probability of animal movement. In the northern half of Canada, one can see areas of high current density around major natural barriers. For example, we observed high current density south of James Bay, an area that has been highlighted as the Ontario-Québec bottleneck^64^. Other natural barriers leading to high current density in northern Canada include Lake Winnipeg in Manitoba, and the McKenzie River and Great Slave Lake in the Northwest Territories. In the south, where natural barriers are further compounded by expanses of built-up areas (e.g., cities) and croplands, areas of high current density tend to be smaller, but are nevertheless present. In fact, some of those correspond to areas already identified by initiatives established to advance connectivity conservation, including Yellowstone to Yukon, Algonquin to Adirondacks, and the Chignecto Isthmus (Figure 2).

### VIGNETTES

We included vignettes of regions with high densities of species at risk because these areas in southern Canada can be difficult to discern on a large-scale, national map, and because they can be overlooked in large-scale connectivity analyses^65^. The resolution of our map is sufficient however, to support land-management decisions and planning at local-scales for many species at risk. By focusing on southern British Columbia (Figure 3a), one can see two prominent north-south corridors of high current density that correspond to the Cascade and Rocky Mountain ranges. In addition, there are regions with diffuse currents – representing areas where many pathways exist for movement - through areas of mostly low movement cost (e.g., areas with ongoing forest management), including adjacent to the Okanagan Valley. More areas of high current density are also apparent in the southern prairies vignette (Figure 3b) than are evident in the national map. Those areas typically follow waterways, such as the Frenchman River westwards from Grasslands National Park, and networks of stream headwaters too hilly for farming to the east. Diffuse currents can be seen across untilled areas with poor soils, typically used as cattle rangelands, including at the Saskatchewan – Alberta border.

Despite a large amount of anthropogenic development, there are still many areas in southern Ontario with high probabilities of movement and therefore opportunities to maintain connectivity (Figure 3c). Pinch points around Lake Simcoe are prominent features in this area, demonstrating the prominence of the corridor along Georgian Bay and leading to the southwest. In southwestern Ontario, many areas of high current density align with ‘back forty’ woodlots that have been maintained by farmers for generations within a landscape of intensive agriculture^66^. Stakeholders and land-use planners across Canada can make use of these current density maps at a 300 m resolution to help inform land use plans about the importance of areas for ecological connectivity.

### METHODOLOGICAL ADVANCES

We used a wall-to-wall, omnidirectional version of circuit theory with nodes placed in buffers outside of the area of interest, allowing for the assessment of connectivity across an entire landscape independent of source and destination nodes^18,20^. We were also fortunate to have access to the input layers for the new Canadian Human Footprint^42^ and a new, more comprehensive national road layer. Those resources allowed us to generate a national cost surface specific for connectivity analyses, with only a few cost categories that we are confident were accurately ranked relative to each other. All of those advances, combined with the use of the Julia implementation of Circuitscape^40^, allowed us to use relatively large tiles (e.g., up to 40 million cells, or approximately 3,600,000 km^2^ and the wall-to-wall method to analyze Canada.

We assessed the sensitivity of current density estimates to cost values and tile sizes following methods similar to those used previously^33,34^ but applied to tiles that were an order of magnitude larger. Consistent with Bowman et al.^34^, we found that current density estimates were relatively insensitive to the absolute cost values assigned to land cover features so long as their rank order was maintained, but that estimates were sensitive to the range of costs used. This held true for both of the tiles we tested, each of which had more than ten million cells, compared to fewer than 500,000 cells tested in the previous study. In addition, our tiles covered more varied landscapes, including the mountainous region of southern British Columbia and the geographically constrained eastern provinces.

When we assessed the sensitivity of current density estimates to tile sizes, we used much larger tiles than previously analyzed (2.25 million cells vs 19,600)^33^ and we used simulated, rather than real landscapes, to control for the effects related to the composition and spatial distribution of cost values. We found that current densities become relatively insensitive to tile sizes above 150,000 km^2^ (> 1.7 million pixels), which fortunately was far smaller than the tile size we could analyze with the Julia implementation of Circuitscape on our computers.

We therefore divided Canada into a relatively small number of tiles (n = 17) that were larger than that threshold. After stitching together the tiles, however, we noticed a few areas with current density anomalies at the seams between tiles. We believe those anomalies resulted from having major geographical barriers on one tile beyond the buffer and overlap of the other. Those barriers likely resulted in current flow being directed to areas that did not receive the same amount of flow on the other tile. We addressed this small number of anomalies by running Circuitscape for five additional tiles, centered and overlapping those regions (Supplemental Figure 6). It is likely that at some point, cloud computing or computing clusters (for example) might be able to facilitate analysis of Canada in its entirety, within a reasonable time frame, without the use of tiles. Doing so could help eliminate seams, although we do not consider the small number of seams in our map to lead to any substantive effects on interpreting patterns of current density.

### VALIDATION OF RESULTS

We validated our current density map using a variety of independently collected wildlife data from across the country. All of the taxa that were analyzed were more likely to be found in areas with high current density compared to nearby areas with lower current density. This was the case for moose and herpetofauna in eastern Canada as well as moose, elk, wolves, and caribou in western Canada. Our results are consistent with those of other studies that have used circuit theory models. For example, current density maps were found to successfully predict areas important for connectivity for wildebeest^67^ (*Connochaetes taurinus*), wild dogs^68^ (*Lycaon pictus*), and elephants^69,70^ (*Loxodonta africana*) in Africa; Tibetan antelope^71^ (*Pantholops hodgsonii*) in China; hedgehogs^72^ (*Erinaceus europaeus*) in Europe; as well as black bear^73^ (*Ursus americanus*), bobcat^74^ (*Lynx rufus*), and lynx^29^ (*Lynx canadensis*) in North America. This is not always the case, however, as the results from some studies that used circuit theory models were either not supported with independent wildlife data^35^ or had mixed results^25,27^.

### COMPARED TO PAST STUDIES

Two recent studies have analyzed connectivity in Canada among protected areas. Barnett and Belotte^36^ modeled an aspirational network of protected areas across North America using both circuit theory and least-cost approaches. Similarly, Brennan et al.^75^ used circuit theory to assess the connectivity of terrestrial protected areas for the entire world, including Canada. These approaches are important for helping to address the goal established by parties to the Convention on Biological Diversity to establish well connected systems of protected areas. However, given that only 12.5% of Canada’s terrestrial areas are currently protected, their approaches could overlook areas important for connectivity in regions with few existing protected areas, including much of southern Canada where most of the country’s species at risk reside. Indeed, this is particularly apparent in southern Ontario where our analysis identifies many areas with high probabilities of movement, while the other studies do not.

Interestingly, there are many similarities among the findings of the three studies. In particular, all three identify the Cascade and Rocky Mountain ranges in the west as being important for connectivity, as well as around the southern tip of James Bay. In contrast, only our study identified the Chignecto Isthmus as an important area, which is the only terrestrial connection between Nova Scotia and the rest of Canada. Our results are also very similar to those from a study focussed on the province of Alberta^10^, which is encouraging because they were able to use a higher resolution cost surface (100m).

### POTENTIAL LIMITATIONS

We acknowledge that decisions made during the course our analysis could have affected the results. We classified large lakes as high cost in contrast to Marrec et al.^10^, who used a medium cost for their analysis that was restricted to the province of Alberta. We chose the high cost because much of the rest of the country includes many very large lakes, which we believe affect the movement of wildlife most of the year. We recognize that this may not be the case in winter, once the lakes have frozen. Similarly, we assigned a high cost to rivers with high flow rates, some of which can freeze in very cold winters. Consequently, patterns of connectivity in winter may differ from our model, suggesting that connectivity may vary seasonally in Canada. We encourage more research on the implications of this idea for land use planning and conservation in Canada.

We are extremely grateful to the organizations and the researchers that made their wildlife data available for our use, but we were unable to obtain wildlife data for the prairies and the relatively pristine north, which would have strengthened the validation of our current density map. We urge all researchers to share their data through portals such as Movebank.org, in order to help advance science and conservation.

Although we used the most current land-cover data available at the time, we recognize that we evaluated a snapshot in time, and so our analysis does not take into consideration the inevitable continued expansion of the human footprint nor the effects of climate change. However, we believe that our current density map, which factors in both natural and human-cause restrictions to movement, accurately identifies existing areas important for connectivity that should be conserved in order to help mitigate the effects of continued development and climate change.

### CONCLUSION

We have produced the first national map of connectivity for Canada that has been validated with independent wildlife data from across the country. Overall, the results of our sensitivity and validation analyses further the findings of previous studies and demonstrate that so long as practitioners accurately rank the cost of land cover types and use large tiles with buffers, current density maps can be used to identify areas important for connectivity for many species, providing confidence for their use in land management decisions. In particular, we believe our map can assist identify important areas that could contribute to national programs such as Parks Canada’s National Corridor Program, the Federal Government’s Two Billion Trees commitment, and for prioritizing areas for restoration and protection of wide-ranging mammal species. Our approach for conducting a nation-wide analysis could be replicated by other countries, potentially with higher-resolution data for smaller countries.

## ACKNOWLEDGEMENTS

We are very grateful to the researchers that made their wildlife telemetry data available through Movebank.org, to Ontario Nature for the herpetofauna road-kill data, and the Government of New Brunswick for the moose road-kill data. This project was undertaken with the financial support of the Government of Canada through the federal department of Environment and Climate Change Canada and the Government of Ontario through the Ministry of Northern Development, Mines, Natural Resources and Forestry.

## Author contributions

R.P. and J.B. conceived the study. R.P. coordinated the project. All authors contributed to data collection; P.O. was the lead for spatial analyses, mapping, and figures. P.O., A.B., and J.B. contributed to statistical analyses. All authors contributed to the writing of the manuscript.

## Additional information

Our input movement cost surface layer and output current density map are available in TIFF format at https://osf.io/z2qs3/; DOI 10.17605/OSF.IO/Z2QS3.

## Corresponding Authors

- Richard Pither, Environment and Climate Change Canada, 1125 Colonel By Drive, Ottawa, K1A 0H3, Ontario, Canada.
- Jeff Bowman, Ontario Ministry of Northern Development, Mines, Natural Resources and Forestry, Peterborough, Ontario, Canada.

