## Supplemental Information for "Areas Important for Ecological Connectivity Throughout Canada"

### SUPPLEMENTARY INFORMATION

Supplementary Table 1. Input Cost Layers. (CHF – Canadian Human Footprint, GHF = Global Human Footprint)

| LAYER | COUNTRY(S) | COST | SOURCE |
|---|---|---|---|
| CHF - Built environments | Canada | 1000 | Agriculture and Agri-Food Canada; Science and Technology Branch 2016 |
| CHF - Croplands | Canada | 100 | Agriculture and Agri-Food Canada; Science and Technology Branch 2016 |
| CHF - Dams | Canada | 1000 | Global Forest Watch Canada 2010 |
| CHF - Forestry (cut between 1985 & 2015) | Canada | 10 | White, J. C., Wulder, M. A., Hermosilla, T., Coops, N. C. & Hobart, G. W. A nationwide annual characterization of 25years of forest disturbance and recovery for Canada using Landsat time series. Remote Sensing of Environment 194, 303–321 (2017). |
| CHF - Mining | Canada | 1000 | Government of Canada; Natural Resources Canada 2017 |
| CHF - Nighttime lights | Canada | 1000 | Annual composite from 2016 generated by NOAA to assess nighttime lights |
| CHF - Oil and gas | Canada | 1000 | Natural Resources Canada 2017 |
| CHF - Pasturelands | Canada | 10 | Agriculture and Agri-Food Canada; Science and Technology Branch 2016 |
| Lakes >= 10ha | Canada | 1000 | Lehner, B., and M. L. Messager. 2016. HydroLAKES Technical Documentation Version 1.0. https://www.hydrosheds.org/page/hydrolakes |
| Rails | Canada | 1000 | Natural Resources Canada's National Railway Network (2012) |
| Roads - minor | Canada | 10 | Poley, L., Schuster, R., Smith, R. & Ray, J. Identifying Differences in Roadless Areas in Canada Based on Global, National, and Regional Road Datasets. In review. |
| Roads - two-lane highway | Canada | 100 | |
| Roads - multi-lane highways | Canada | 1000 | |
| Elevation > 2300m | Canada & U.S. | 1000 | Global Multi-resolution Terrain Elevation Dataset, USGS 2010 |
| Glaciers | Canada & U.S. | 1000 | CanVec Series, Hydrographic Features 2017 |
| Ocean | Canada & U.S. | 1000 | NRCan Atlas of Canada Data (2017) |
| Rivers > 28m3/sec | Canada & U.S. | 1000 | HydroRIVERS : Lehner, B. & Grill, G. Global river hydrography and network routing: baseline data and new approaches to study the world's large river systems. Hydrological Processes 27, 2171–2186 (2013). |

| | | | |
|---|---|---|---|
| Sea Ice | Canada & U.S. | 10 | USGS North America Glaciers and Sea Ice |
| Slopes > 30 degrees | Canada & U.S. | 1000 | Global Multi-resolution Terrain Elevation Dataset, USGS 2010 |
| GHF - Built environments | U.S. | 1000 | Venter, O. et al. Sixteen years of change in the global terrestrial human footprint and implications for biodiversity conservation. Nat Commun 7, 12558 (2016). |
| GHF - Croplands | U.S. | 100 | |
| GHF - Nightime lights | U.S. | 1000 | |
| GHF - Pasturelands | U.S. | 10 | |
| GHF - Rails | U.S. | 1000 | |
| GHF - Road buffer (500 - 1000m) | U.S. | 1 | |
| GHF - Road buffer (500m) | U.S. | 100 | |
| GHF - Roads | U.S. | 1000 | |

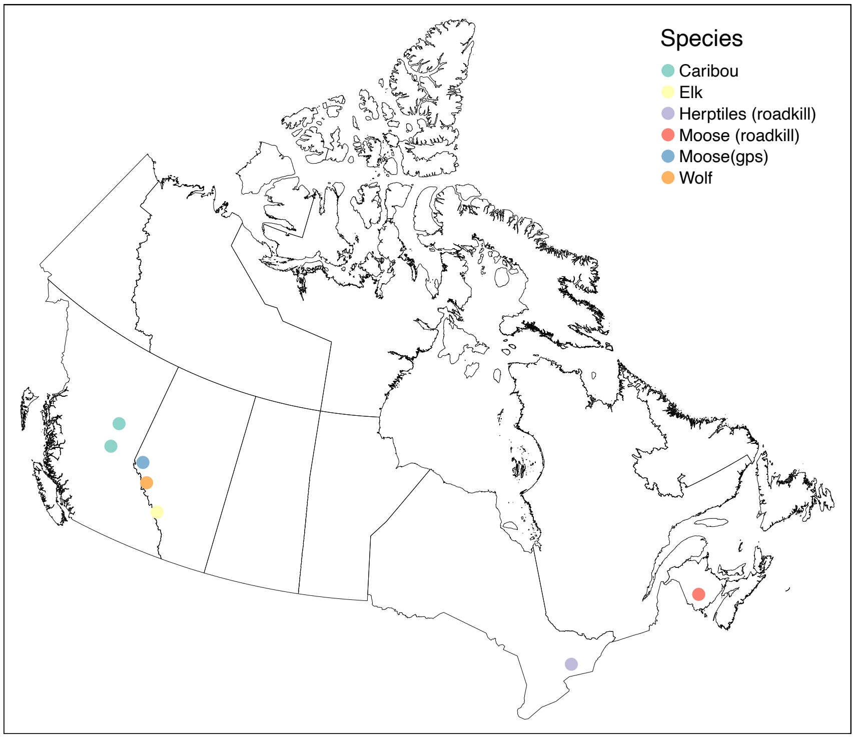


Supplementary Figure 1. Locations of independent wildlife data: GPS collar data for caribou in British Columbia, moose, wolf, and elk in Alberta; herptofauna road kill in Ontario, and moose road kill in New Brunswick.

Supplementary Table 2. Cost scenarios used to compare across two landscapes with four cost categories.

| **Scenario** | **Low** | **Medium Low** | **Medium High** | **High** | **Range** |
|---|---|---|---|---|---|
| C1 | 0.1 | 0.5 | 1 | 1.5 | 1.4 |
| C2 | 1 | 1.5 | 2.25 | 3.375 | 2.375 |
| C3 | 1 | 1.5 | 2.25 | 225 | 224 |

| | | | | | |
|---|---|---|---|---|---|
| C4 | 1 | 2 | 3 | 4 | 3 |
| C5 | 1 | 2 | 3 | 300 | 299 |
| C6 | 1 | 5 | 7.5 | 10 | 9 |
| C7 | 1 | 5 | 7.5 | 750 | 649 |
| C8 | 1 | 100 | 150 | 200 | 199 |
| C9 | 1 | 10 | 100 | 1000 | 999 |
| C10 | 10 | 100 | 1000 | 10000 | 9990 |

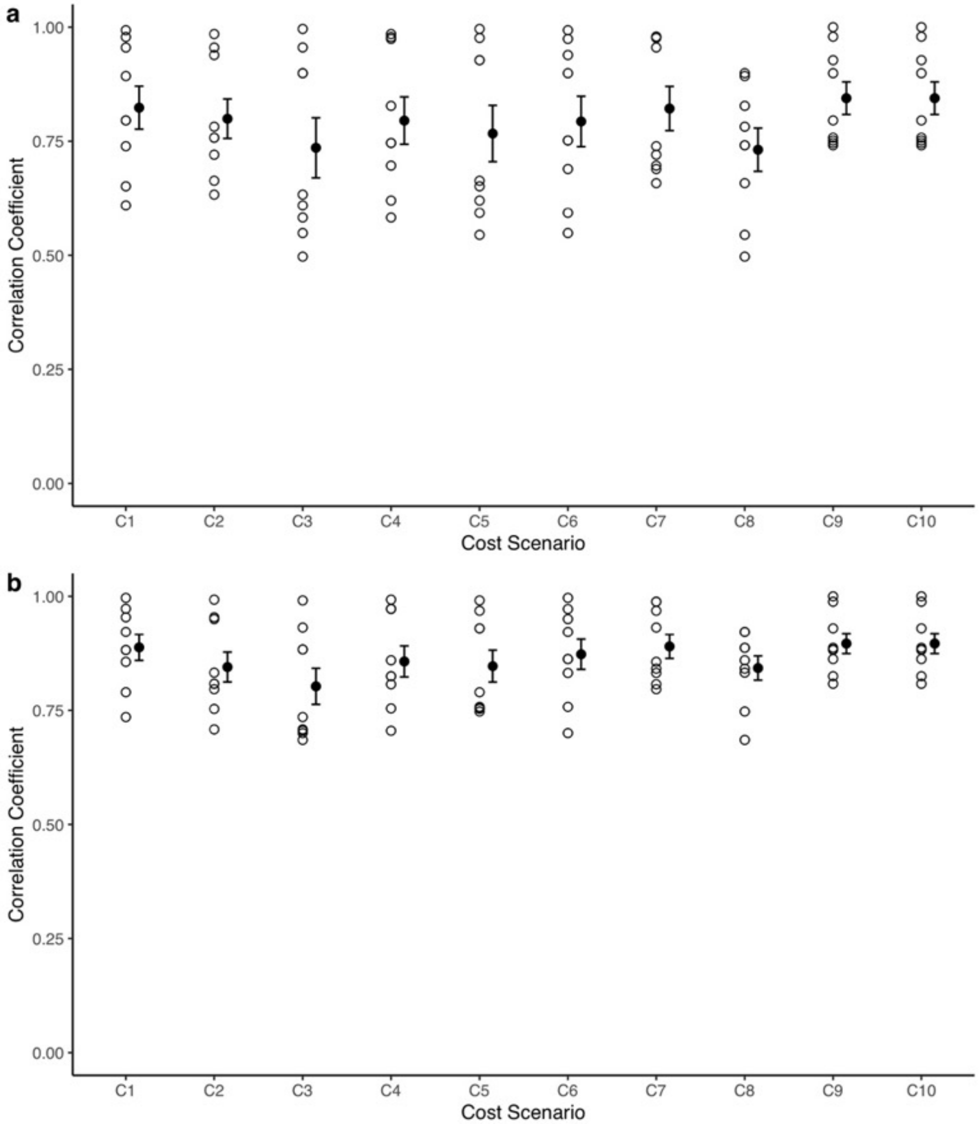


Supplementary Figure 2. Correlation of mean current densities among 10 cost value scenarios for **a)** east coast provinces and **b)** southern British Columbia. Scenarios 9 and 10 had highest mean correlation values with other scenarios (solid circles denote scenario means, and bars +/- one standard error).

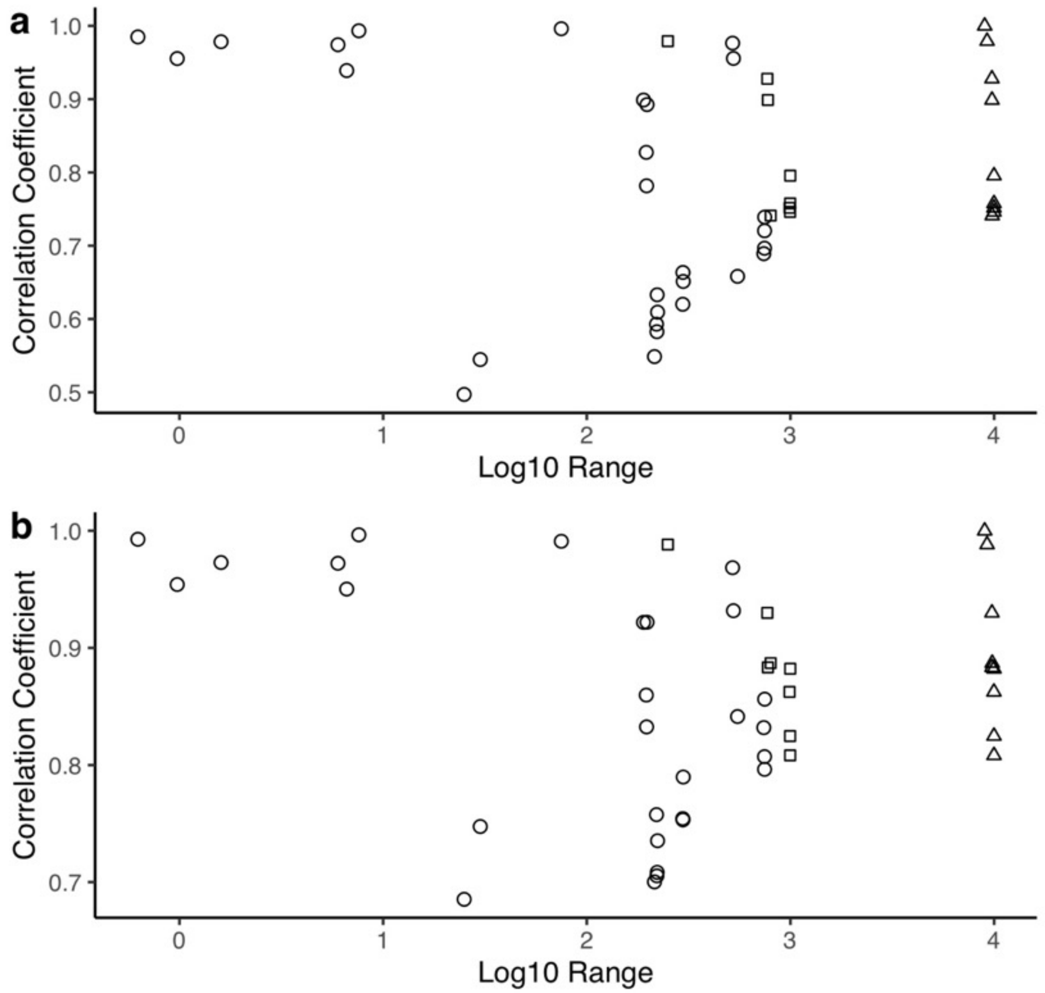


Supplementary Figure 3. Effect of the range of cost values on correlation of current density across landscapes. This figure shows the effect of the absolute difference in the range of cost values between pairs of landscapes (log10-transformed) on the Spearman rank correlation between current density estimates in pairs of landscapes for two study area tiles: **a)** east coast provinces and **b)** southern British Columbia. Scenarios #9 & #10 identified by square and triangle symbols, respectively.

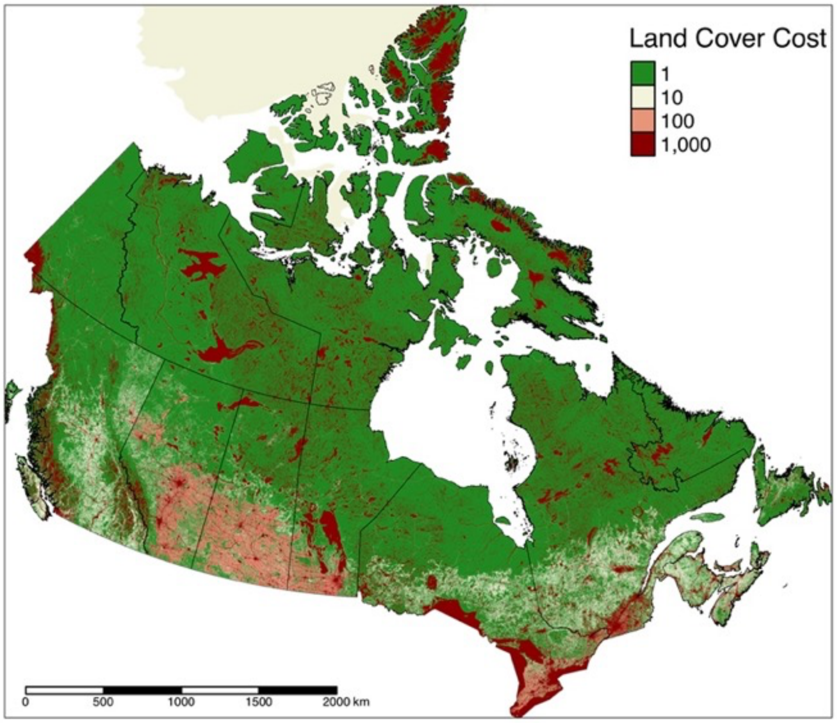

Supplementary Figure 4. Cost surface map of Canada, with four movement costs.

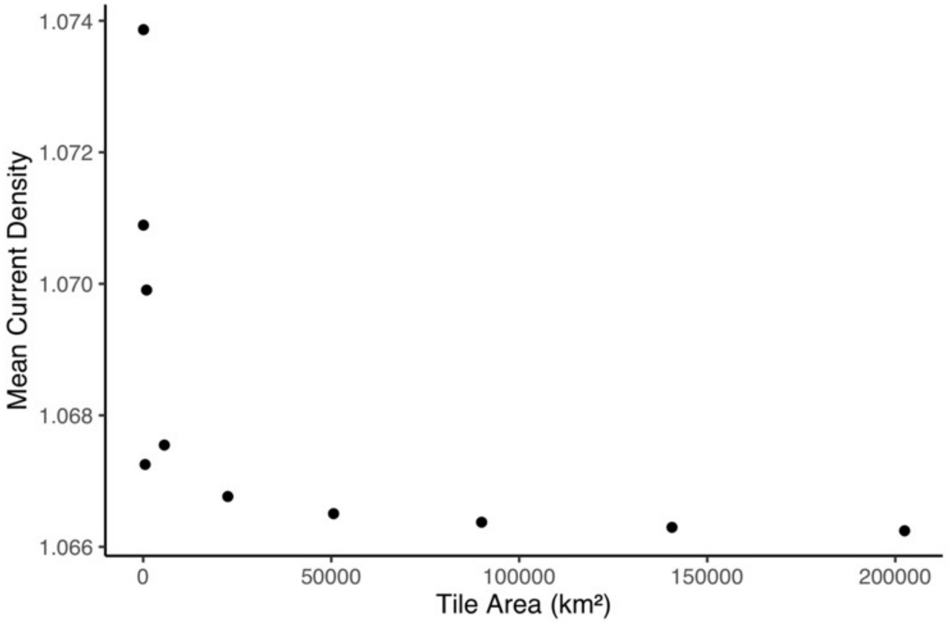

Supplementary Figure 5. Effect of tile size on mean current density. Analysis was conducted on simulated but identical landscapes, to control for composition and spatial distribution of cost values. The same pattern was found for the minimum, maximum, and standard deviation of current densities.

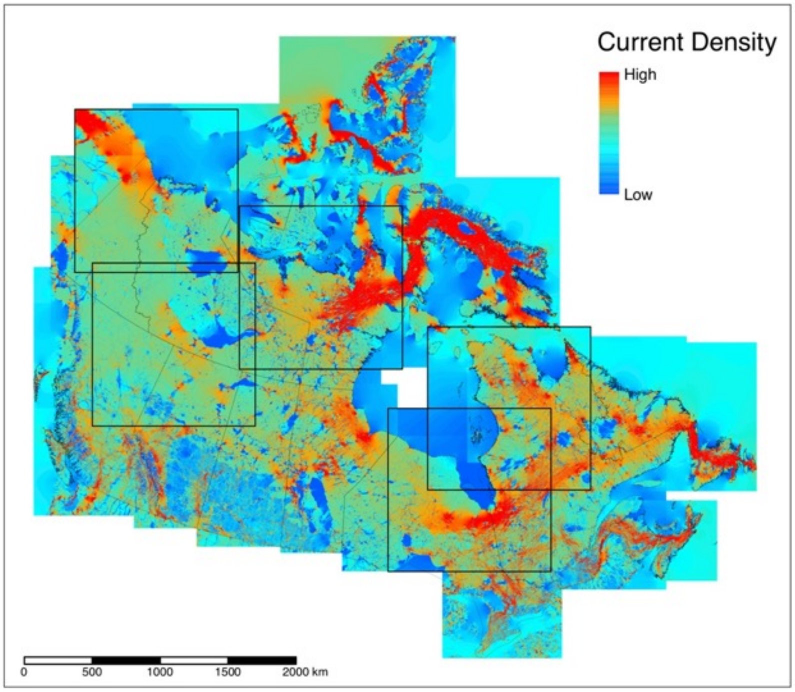

Supplementary Figure 6. Map showing all 22 tiles used for the analysis over raw current density. Black boxes indicate the 5 tiles that were required to address anomalies at the seams.
